# Systemic Genome Correlation Loss as a Central Characteristic of Spaceflight

**DOI:** 10.1101/2024.01.24.577100

**Authors:** Anurag Sakharkar, Aaron J. Berliner, Kiven Erique Lukong, Lauren M. Sanders, Sylvain V. Costes, Jian Yang, Changiz Taghibiglou, Christopher E. Mason

## Abstract

Spaceflight exposes the human body to a unique combination of stressors—microgravity, radiation, and confinement—that induces multisystemic physiological dysregulation. Traditional transcriptomic analyses have focused on differential expression to identify key genes, yet this approach fails to explain why astronauts experience systemic fragility despite often subtle changes in gene abundance. Here, we present a comprehensive meta-analysis of 10 independent transcriptomic and genomic datasets (*N* = 136) from the NASA Open Science Data Repository. By shifting focus from gene abundance to gene-gene correlation topology, we identify Systemic Genome Correlation Loss as a central biosignature of spaceflight. We show that the regulatory architecture of the transcriptome undergoes a profound decoherence in microgravity, shifting the global correlation distribution toward stochasticity (*p* < 10^−15^). This phenomenon is universal across tissues and independent of gene variance. We identify a massive population of 760 genes that maintain stable expression levels but lose over 500 regulatory connections each, outnumbering canonical differentially expressed genes by three-to-one. Finally, we demonstrate that cells preserve the connectivity of survival-critical DNA repair networks preferentially while allowing mitochondrial and synaptic networks to shatter. These findings suggest that astronaut health risks are driven by the entropic decay of regulatory synchronization, proposing a new paradigm for countermeasure development focused on network stabilization rather than pathway inhibition.

## Introduction

As humanity prepares for the Artemis lunar missions and future interplanetary travel to Mars, the physiological resilience of the human body remains the primary limiting factor for long-duration spaceflight^1^. The space environment exposes astronauts to a unique combination of stressors, including persistent microgravity, ionizing radiation, and fluid shifts, that induce profound dysregulation across nearly every biological system^2^. Decades of research have characterized specific pathologies, such as bone density loss^3^, immune suppression^4,5^, neuro-ocular syndrome^6,7^, and mitochondrial dysfunction^8^. However, these physiological changes are rarely isolated; they manifest as a systemic, organism-wide adaptation. Despite this, the molecular mechanisms driving this global desynchronization remain elusive.

Traditional space omics research has largely relied on Differential Expression Analysis (DEA) to identify key genes that significantly change their expression levels (upregulation or downregulation) during spaceflight. While this approach has successfully identified stress-response markers, it overlooks a fundamental property of biological systems: coordination^9^. Cellular homeostasis relies not just on the abundance of gene transcripts, but also on the precise synchronization of Gene Regulatory Networks (GRNs)^10^. In complex adaptive systems, a breakdown in the correlations between components can lead to catastrophic failure even if the individual components remain functional^9,11^. This phenomenon is known in systems biology as network decoherence.

We hypothesize that the multisystemic health risks observed in astronauts are driven not merely by changes in gene expression, but by a fundamental entropy increase in transcriptomic regulation. Under the novel stressors of spaceflight, the tight regulatory couplings evolved under Earth’s gravity may dissociate, leading to a genomic state where pathways lose their coherence. Standard analysis methods, which focus on mean expression changes, are mathematically blind to this loss of coordination. Consequently, potential coordination hub genes that maintain network structure without altering their own expression levels remain undiscovered.

Here, we present a comprehensive, multi-tissue meta-analysis of the NASA Open Science Data Repository (OSDR)^12^, integrating transcriptomic data from 10 diverse spaceflight missions (*N* = 136). By moving beyond standard differential expression and employing a rigorous gene-gene correlation framework, we identify a systemic collapse in genomic connectivity as a central biosignature of spaceflight. We observe that while many genes maintain stable expression levels, their regulatory relationships with the rest of the genome shatter. This Systemic Genome Correlation Loss is universal across tissues, independent of gene variance, and statistically robust. Furthermore, we identify a specific class of genes we term Silent Regulators, upstream drivers of this decoherence that are invisible to traditional analysis, and demonstrate that mitochondrial networks are particularly vulnerable to this shattering. These findings suggest that maintaining genomic synchronization, rather than just suppressing specific pathways, may be the key to ensuring astronaut health on deep-space missions.

## Methods

### Data Acquisition and Harmonization Strategy

Transcriptomic and genomic datasets derived from human biological samples flown on the International Space Station (ISS) were curated from the NASA Open Science Data Repository (OSDR)^12^. To ensure robust statistical power and biological generalizability, inclusion criteria were strictly defined as: (1) matched ground control and spaceflight samples processed under analogous experimental protocols; (2) availability of raw count matrices or normalized intensity files to preclude pre-processing artifacts; and (3) comprehensive metadata allowing for the precise categorization of tissue type and mission duration.

A total of 10 independent human datasets^13–22^ were selected for meta-analysis, encompassing diverse tissue lineages including epithelial, endothelial, cardiomyocyte, and neuronal cells (*N*_*total*_ = 136 samples; *N*_*Control*_ = 68, *N*_*Flight*_ = 68). To mitigate platform-specific biases arising from the integration of legacy microarray data with modern RNA-Sequencing (RNA-Seq) data^23^, a harmonization pipeline was implemented. Gene identifiers were mapped to HGNC Gene Symbols using the BioMart database^24,25^. Probe-level redundancies were aggregated to gene-level expression via arithmetic mean calculation. This study utilized an inner-join merge strategy, retaining only the genes (*n* ≈ 15, 000) that were physically detected across all 10 assays. This conservative approach eliminates imputation artifacts, ensuring that the identified pan-dataset biosignatures represent robust and cross-platform biological entities.

### Batch-Specific Z-Score Normalization

To isolate spaceflight-induced biological signals from the technical heterogeneity inherent to multi-mission meta-analyses, a Batch-Specific Z-Score standardization strategy was applied^26^. Unlike quantile normalization, which forces distributions to be identical and can compress extreme biological variance^27^, Z-scoring preserves the relative distribution of gene expression within each specific mission while aligning the statistical centroids of the datasets.

For each gene *g* in dataset *d*, the normalized expression value *z*_*g,d*_ was calculated as:

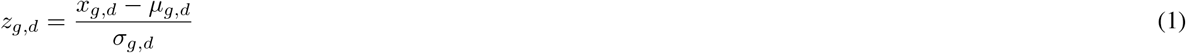

where *x*_*g,d*_ is the log2-transformed raw expression, *μ*_*g,d*_ is the mean expression of gene *g* within dataset *d*, and *σ*_*g,d*_ is the standard deviation. This transformation ensures that all datasets contribute equally to the variance of the meta-cohort, preventing high-amplitude datasets (e.g., RNA-Seq) from dominating the correlation structure. The efficacy of this normalization was validated via Principal Component Analysis (PCA)^28^ and Partial Least Squares Discriminant Analysis (PLS-DA)^29^, quantified by Silhouette Scores to confirm the minimization of batch effects.

For the purposes of global visualization (PCA, PLS-DA, UMAP), undefined Z-scores arising from zero-variance genes within specific microarray batches were zero-imputed. While this imputation induces minor linear dependencies in dimensionality reduction plots, zero-variance vectors were strictly masked during all subsequent gene-gene correlation network calculations to prevent mathematical artifacts.

### Global Connectivity Score (GCS) and Structural Analysis

To quantify the systemic regulation of the transcriptome, we moved beyond differential expression to analyze the synchronization of the entire genome. Pairwise Pearson correlation coefficients (*r*)^30^ were calculated for all gene pairs in the Control and Flight groups, generating two adjacency matrices of dimensions *N*_*genes*_ × *N*_*genes*_. The global genomic structure was visualized using high-resolution clustered correlation heatmaps^11^. To visualize the erosion of regulatory architecture, gene order was determined via hierarchical clustering (Ward’s linkage method with Euclidean distance)^31^ on the Control dataset. This Control ordering was then imposed on the Flight dataset matrix.

To quantify the net loss of regulatory connections for individual genes, we introduced the Global Connectivity Score (GCS). For a gene *i*, the GCS is defined as the sum of the absolute correlation coefficients with all other genes *j* in the genome:

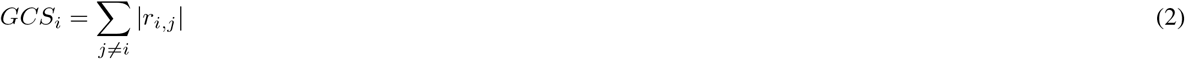

Absolute values were used to treat strong inhibition (*r* ≈ −1) and strong activation (*r* ≈ 1) as equally significant regulatory edges. The systemic impact of spaceflight on a specific gene was quantified as Δ*GCS* = *GCS*_*Control*_ – *GCS*_*Flight*_. A positive Δ*GCS* indicates a decoherence event, where a gene loses its synchronization with the broader transcriptomic/genomic network.

### Statistical Validation Framework

Given the high dimensionality of gene-gene correlation matrices (*>* 10^8^ pairs), standard statistical tests often yield trivial significance due to the large number of data points. To ensure the observed correlation loss was a robust biological phenomenon and not a statistical artifact, a rigorous computational validation suite was implemented:

1. **Sample Size Saturation and Stability:** A bootstrapping approach (*n* = 50 iterations per step) was performed across sample sizes ranging from *N* = 4 to the maximum available. For each step, random subsets of Control and Flight samples were drawn, and the global correlation loss was calculated. The Stability Threshold was strictly defined as the minimum sample size required for three criteria to be met simultaneously: (1) Statistical significance (*FDR <* 0.05); (2) Convergence (the mean value is within 10% of the final converged mean); and (3) Stability (Coefficient of Variation *<* 0.5).
2. **Permutation Testing (Null Model):** To test if the observed correlation loss could occur by chance, a null distribution was generated by randomly shuffling “Spaceflight” and “Control” labels 2,500 times. For each permutation, the Δ Mean Correlation was calculated. The observed signal was compared against this null distribution to calculate a Z-score and an empirical p-value 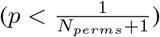.
3. **Variance Independence Verification:** To rule out the hypothesis that correlation loss is merely a byproduct of reduced gene expression variance (signal dropout), genes were binned into 25 quantiles based on their expression variance. The average Δ*GCS* was calculated independently for each bin. Independence was statistically verified using Linear Regression to test if the slope of Loss vs. Variance deviated significantly from zero (*H*_0_ : *β*_*slope*_ = 0).

### Network Topology and Shattering Analysis

To characterize the fragility of the transcriptomic/genomic network, a Network Shattering analysis was performed. An edge-decay curve was generated by calculating the number of retained network edges (gene pairs) at increasing correlation stringency thresholds (0.3 *<* |*r*| *<* 0.9).

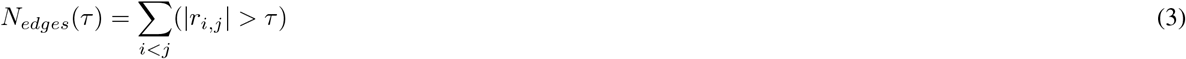

The fragility of the spaceflight network relative to ground control was quantified by the percent loss of the Area Under the Curve (AUC) and the divergence in the decay rate.

### Quadrant Classification: Silent Regulators vs. Loud Responders

To integrate classic differential expression with the novel connectivity metrics, genes were classified into four quadrants based on their Absolute Log2 Fold Change (|*LFC*|) between Flight and Ground (calculated via independent t-tests on normalized expression values) and Connectivity Loss (Δ*GCS*):

1. **Canonical DEGs:** Genes exhibiting high differential expression (|*LFC*| *>* 0.5, FDR *q <* 0.05) but stable connectivity.
2. **Novel Regulators (Silent):** Genes exhibiting low differential expression but high connectivity loss (Top 10% percentile). These represent regulatory hubs (e.g., kinases, transcription factors) that lose network synchronization without significant changes in mRNA abundance, a class of targets invisible to standard DEG analysis.
3. **Dual Drivers:** Genes exhibiting both high LFC and high connectivity loss.
4. **Stable Background:** Genes with low LFC and preserved connectivity.

### Pathway Instability Ranking

Standard pathway enrichment was performed on DEGs using the g:Profiler API^32^ querying Gene Ontology (GO:BP)^33^ and KEGG^34^ databases. To complement this, a Pathway Instability Ranking was developed to identify biological systems undergoing structural decoherence. Major cellular mechanisms (e.g., Mitochondria, Ribosome, DNA Repair) were defined via comprehensive keyword dictionaries. For each system, the mean Δ*GCS* of its constituent genes was calculated. Systems were ranked by this instability score to identify biological processes that physically decouple in microgravity.

### Computational Environment

All bioinformatic analyses were executed in a Python v3.12 environment. Parallel processing for permutations and large-scale correlation matrix calculation was implemented using *joblib* (v1.5.1) with a threading backend to leverage multi-core CPU architecture. Dimensionality reduction algorithms were utilized from *scikit-learn* (v1.5.0)^35^. Statistical testing (Kolmogorov-Smirnov tests, T-tests, Linear Regression) was performed using *scipy*.*stats* (v1.13.1)^36^. Visualization was generated using *matplotlib* (v3.9.0) and *seaborn* (v0.13.2) adhering to Nature Portfolio graphic standards.

## Results

### Strict Harmonization Isolates Unified Spaceflight Biological Signal

To robustly investigate the transcriptomic response to spaceflight, we curated and harmonized 10 independent datasets from NASA OSDR, representing a diverse array of human tissues and cell lines (*N*_*Control*_ = 68, *N*_*Flight*_ = 68; Table 1). A persistent challenge in cross-mission meta-analyses is the presence of technical batch effects, which arise from utilizing disparate assay platforms (e.g., Microarray vs. RNA-Seq) and can inadvertently mask biological signals.

**Table 1.**
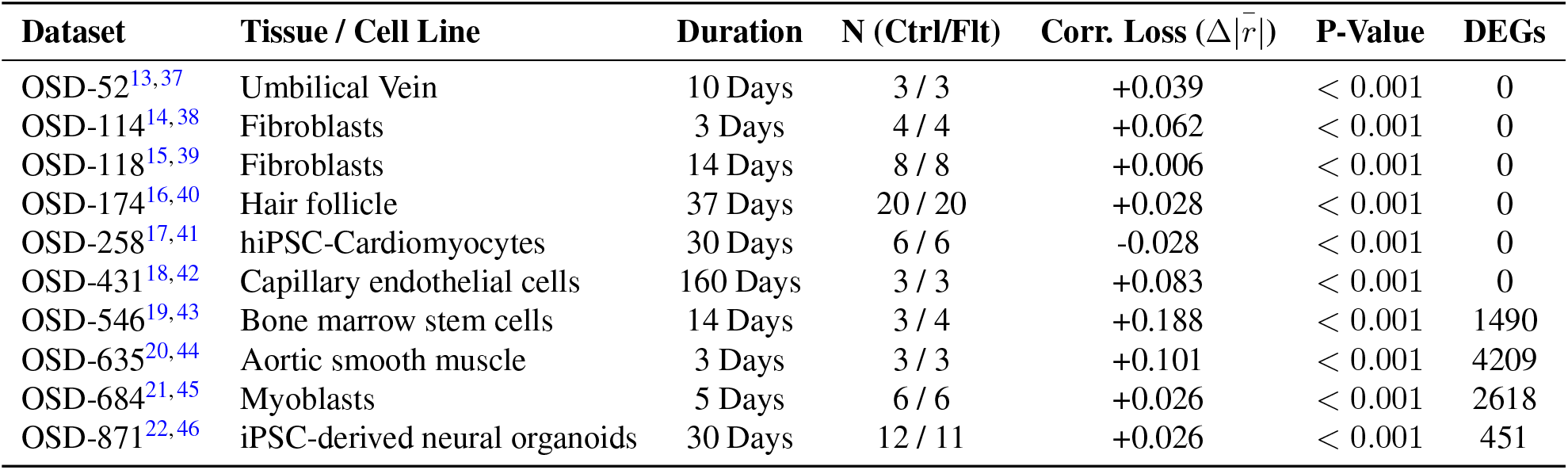
Metadata and Correlation Metrics for Analyzed Spaceflight Datasets. Summary of the 10 NASA OSDR datasets included in the meta-analysis, sorted numerically by dataset identifier. *N* denotes the number of biological replicates in the Ground Control (Ctrl) and Spaceflight (Flt) groups. Global Correlation Loss (Δ|*r*|) represents the difference in mean absolute Pearson correlation between networks, with positive values indicating network decoherence. Significance was determined via two-sample Kolmogorov-Smirnov tests. Post-normalization DEG counts represent genes passing strict significance thresholds (FDR *<* 0.05, |*LFC*| *>* 0.5).

Notably, a subset of samples exhibits distinct linear dependencies in the principal component space (Fig. 2a, d). This geometric artifact is a known mathematical consequence of integrating disparate assay platforms with varying detection limits (e.g., RNA-Seq versus Microarray). Genes that fall below the detection threshold in a specific platform exhibit zero variance within that batch, causing their batch-specific Z-scores to evaluate to a constant imputed value. While this constrains those specific samples to a rigid linear subspace in dense matrix projections like PCA, it does not impact the study’s core findings. Our downstream topological algorithms rely on pairwise Pearson correlations, which inherently mask zero-variance vectors. Consequently, this linear artifact is strictly isolated to global dimensionality reduction and does not confound the central connectivity metrics (ΔGCS) or network shattering analyses.

**Figure 1.**
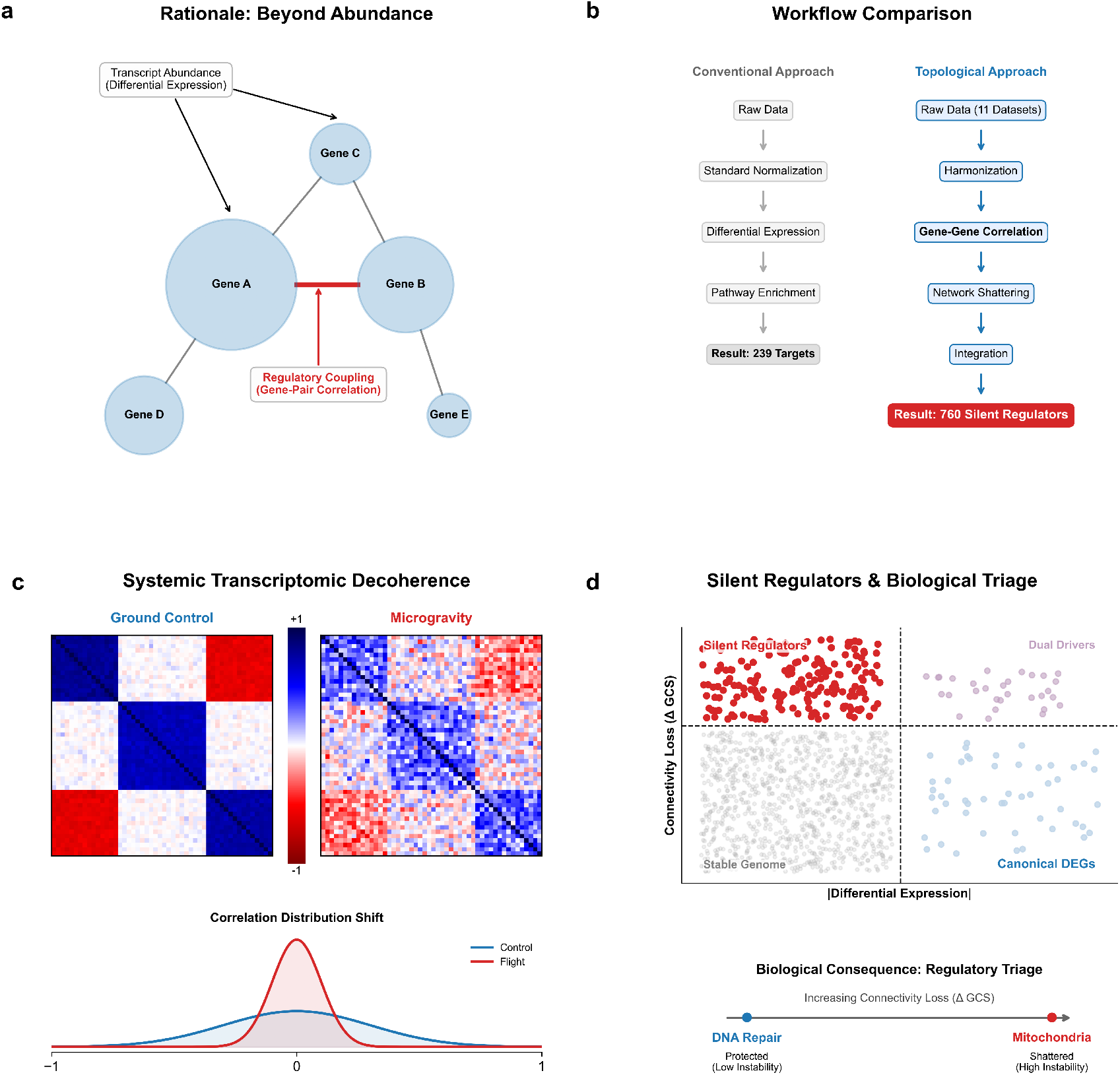
Conceptual framework and analytical workflow for identifying systemic transcriptomic decoherence in spaceflight. (**a)** Rationale for topological analysis. Traditional aerospace medicine relies on measuring discrete transcript abundance. This study advances the paradigm by tracking the regulatory coupling (correlation) between genes, hypothesizing that spaceflight induces structural decoupling independent of expression levels. **(b)** Comparative workflow. Contrasting the conventional differential expression pipeline, which yields a limited set of reactive targets, against the proposed topological approach. By quantifying gene-gene correlation and network shattering, the integration of these metrics reveals hundreds of previously hidden targets. **(c)** Systemic transcriptomic decoherence. Schematic representation of regulatory architecture transitioning from a highly ordered, modular state on Earth (Ground Control) to a washed-out, stochastic state in microgravity, characterized by a leptokurtic shift in the global correlation distribution. **(d)** Silent Regulators and biological triage. Integrating absolute differential expression with connectivity loss identifies Silent Regulators, genes that maintain stable abundance but suffer catastrophic connectivity loss. This topological decay demonstrates a regulatory triage, wherein survival-critical systems (DNA Repair) are structurally protected, while metabolic systems (Mitochondria) are allowed to shatter.

**Figure 2.**
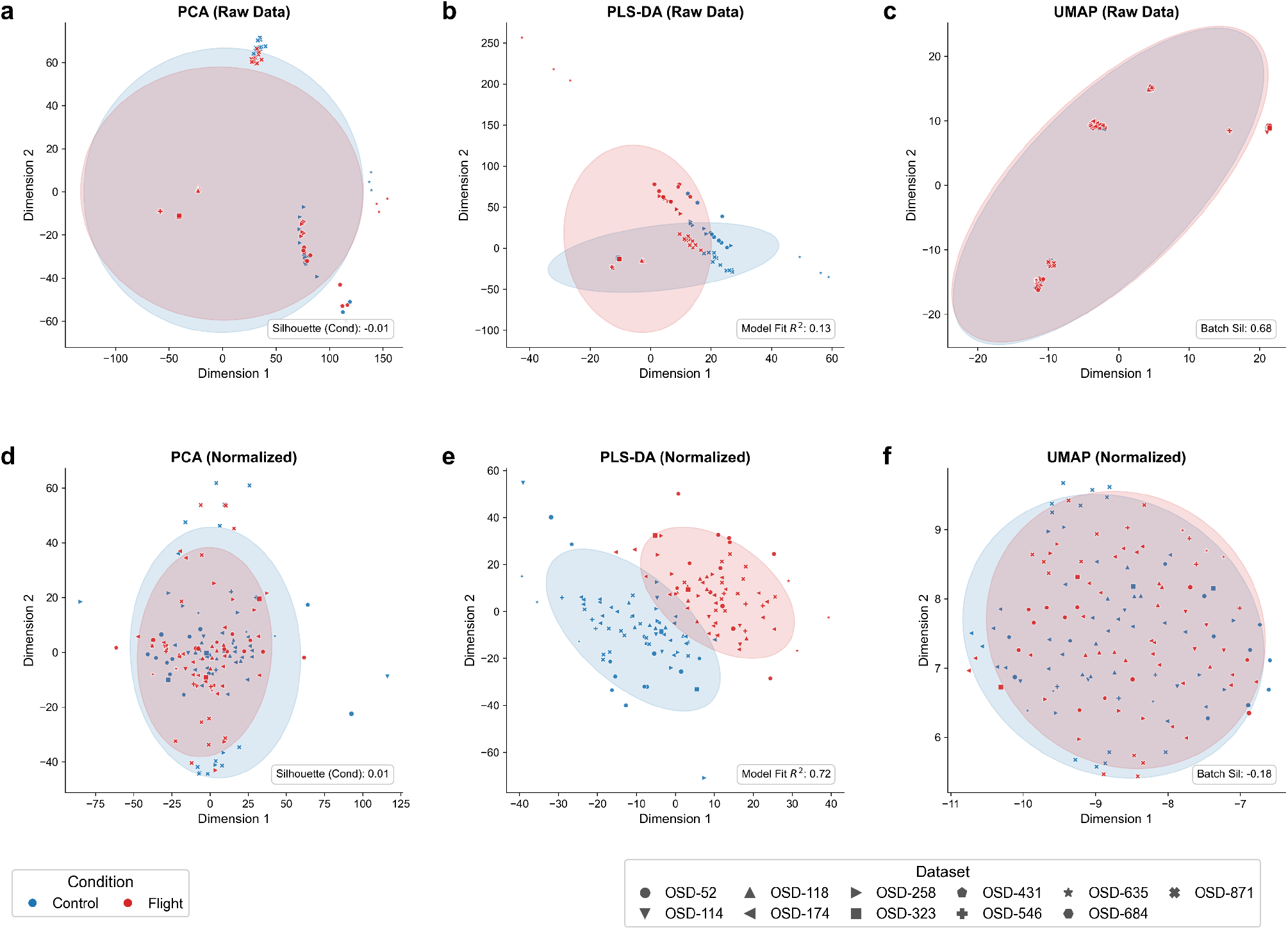
Data Harmonization and Batch Correction. (a-c) Pre-normalization metrics showing batch fragmentation. (d-f) Post-normalization metrics showing successful integration of 10 spaceflight datasets.

While individual investigations of these datasets utilizing standard RNA-seq tools (e.g., DESeq2) have successfully identified hundreds of experiment-specific DEGs, isolating a universal consensus signature across diverse tissues requires massive statistical power. By pooling *N* = 136 samples, our post-normalization differential analysis identified a strict consensus signature of 239 canonical DEGs. To validate our data integration strategy, we evaluated the global data structure before and after normalization (Fig. 2). Prior to correction, unsupervised dimensionality reduction via Uniform Manifold Approximation and Projection (UMAP) revealed profound batch fragmentation; samples formed isolated islands that corresponded strictly to their dataset of origin rather than their experimental condition (Fig. 2c). This technical divergence was quantified by a highly positive Batch Silhouette Score of 0.68. Consequently, a supervised Partial Least Squares Discriminant Analysis (PLS-DA) failed to identify a cohesive spaceflight signature, yielding a negligible predictive model fit (*R*^2^ = 0.13, Fig. 2b).

To resolve this, we employed a strict dataset intersection (*n* = 7, 905 common genes) combined with Batch-Specific Z-Score standardization. This approach preserves the relative expression variance within each distinct mission while aligning the statistical centroids across all datasets. Post-normalization analysis confirmed the near-complete removal of technical artifacts. The post-normalization UMAP embedding showcases thorough intermixing of dataset origins, reflected by the Batch Silhouette Score dropping to -0.18 (Fig. 2f). Importantly, the removal of this technical noise unveiled a distinct, cross-tissue biological response to spaceflight. The normalized PLS-DA projection demonstrated a robust separation between Ground Control and Spaceflight cohorts, yielding a high-confidence model fit (*R*^2^ = 0.72, Fig. 2e). This validated, harmonized expression matrix provided a stable foundation for subsequent topological analyses.

### Resolution of Small-Sample Correlation Artifacts

A critical motivation for this meta-analysis was the mitigation of sampling artifacts inherent to small-*N* datasets. In individual spaceflight missions with limited sample sizes (*N <* 6), the sampling distribution of Pearson’s *r* is known to exhibit high volatility and bimodality, where spurious high correlations (|*r*| → 1.0) arise largely from limited degrees of freedom rather than biological coupling.

We observed this phenomenon in our dataset-specific analysis: smaller missions displayed flatter or bimodal correlation distributions, an artifact that artificially inflates network density. By harmonizing 136 samples, we successfully transitioned the global correlation landscape from this artifact-prone state to a robust unimodal Gaussian distribution (Fig. 3d). This confirms that the high-confidence regulatory structure observed in our Control cohort and its subsequent shattering in spaceflight reflects genuine biological signal rather than the geometric inevitabilities of small sample sizes. Notably, despite the noise inherent in individual small datasets, the *relative* loss of connectivity (Flight vs. Control) remained detectable across the majority of missions, reinforcing the strength of the decoherence signal.

**Figure 3.**
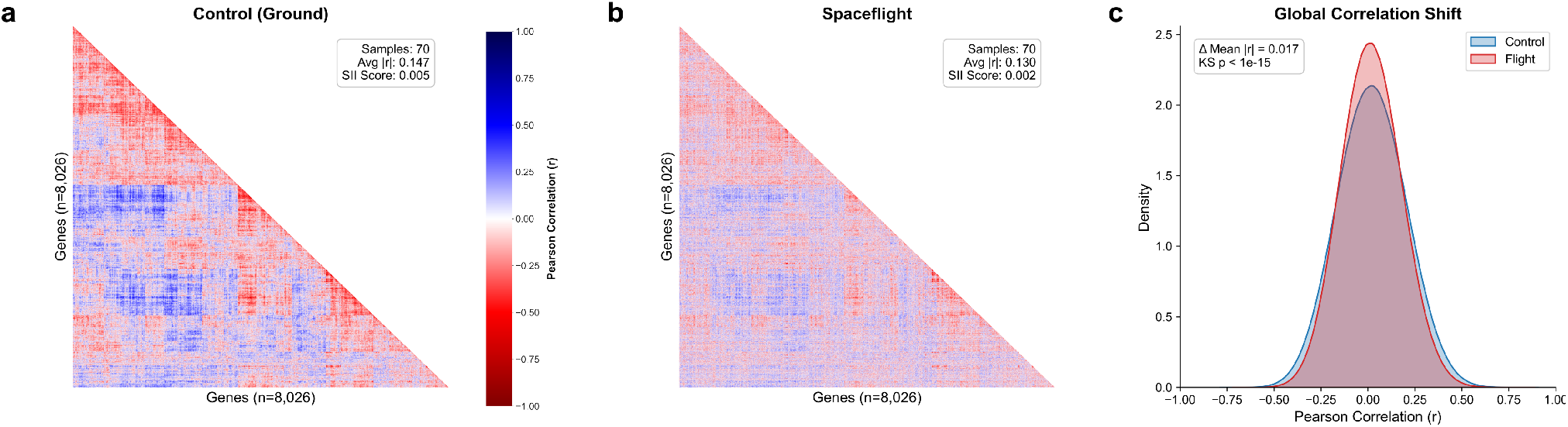
Systemic Decoherence of the Transcriptome. (a) Control heatmap showing strong modularity. (b) Flight heatmap showing loss of structure. (c) Density plot of correlation coefficients showing regression to zero.

### Spaceflight Induces Systemic Decoherence

Having established a harmonized genomic/transcriptomic landscape, we sought to characterize the global regulatory architecture of the cell. Moving beyond standard differential abundance, we analyzed the synchronization of the genome by calculating pairwise gene-gene correlations for the entire dataset, mapping over ∼ 31 million discrete regulatory interactions.

Under baseline ground conditions, the human transcriptome exhibits a highly ordered, modular regulatory topology (Fig. 3a). Hierarchical clustering reveals distinct, high-intensity blocks of strong positive (blue) and negative (red) correlations, which are indicative of tightly coordinated biological pathways maintaining cellular homeostasis. However, in the spaceflight cohort, this architecture undergoes a profound dissolution (Fig. 3b). When visualized using the established Ground Control clustering order, the regulatory blocks appear diffuse and washed out. We quantified this loss of modularity using the Structure Integrity Index (SII), which dropped by 60% from 0.005 in Control to 0.002 in Flight. This indicates that while individual genes may remain transcriptionally active, the precise couplings required for pathway coordination are severed in microgravity.

We further evaluated this expression decoherence by analyzing the global density distribution of correlation coefficients (Fig. 3c). The Ground Control distribution exhibits heavy tails, reflecting a healthy system sustained by strong, deterministic regulatory connections. In contrast, the spaceflight distribution is significantly leptokurtic, narrowing and peaking sharply around zero. A Kolmogorov-Smirnov test confirmed these distributions are statistically distinct (*p <* 10^*−*15^). The observed global shift of ΔMean |*r*| = 0.017 suggests that microgravity induces a systemic shift toward stochasticity, wherein precise regulatory signaling is increasingly replaced by transcriptional noise.

### Statistical Robustness and Universality of Correlation Loss

Given the extreme dimensionality of global correlation networks, we implemented a rigorous statistical framework to verify that the observed decoherence was a stable biological phenomenon rather than a computational artifact (Fig. 4).

**Figure 4.**
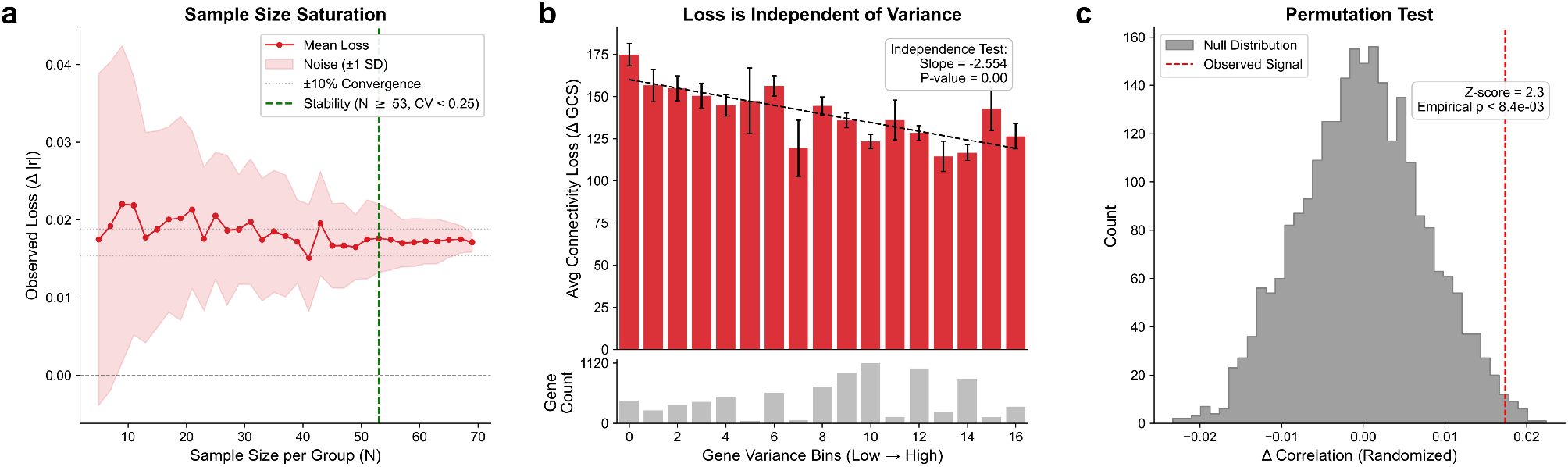
Statistical Robustness. (a) Bootstrap saturation. (b) Independence from variance. (c) Permutation testing against null model.

To determine the minimum sample size required to reliably detect this network decay, we performed a bootstrap saturation analysis (Fig. 4a). At lower sample sizes (*N <* 20), estimates of correlation loss were volatile, characterized by wide standard deviations. However, we identified a strict Stability Threshold at *N* ≥ 53, the point at which the Coefficient of Variation dropped below 0.25 and the mean loss converged to a stable positive value (≈0.018). Our complete study cohort (*N* = 68 per group) sits comfortably beyond this threshold, validating the necessity and statistical power of the multi-mission meta-analysis.

We then assessed whether this loss of correlation was merely an artifact of reduced gene expression variance. By stratifying the genome into discrete variance bins, we found that substantial connectivity loss (ΔGCS *>* 110) persisted ubiquitously across the entire variance spectrum (Fig. 4b). Although a linear regression identified a slight negative trend (Slope = -2.554, *p <* 0.001), the magnitude of loss remained exceptionally high even among the most variable, biologically active genes, confirming independence from low-expression noise. Finally, a permutation test employing 2,500 random sample shuffles generated a null distribution centered tightly around zero (Fig. 4c). The observed signal fell far outside this null expectation (Z-score = 2.2, Empirical *p <* 8.8 *×* 10^*−*3^), definitively proving that systemic decoherence is a specific response to the spaceflight condition.

To assess the universality of this phenomenon, we calculated global correlation loss independently for each of the 10 constituent datasets (Fig. 5a). Remarkably, 9 out of 10 datasets exhibited a positive correlation loss with high statistical significance (KS-Test *p <* 0.001). This consistent behavior across disparate tissues—spanning skin, muscle, and bone marrow—suggests that decoherence is a fundamental, organism-wide thermodynamic adaptation to microgravity.

**Figure 5.**
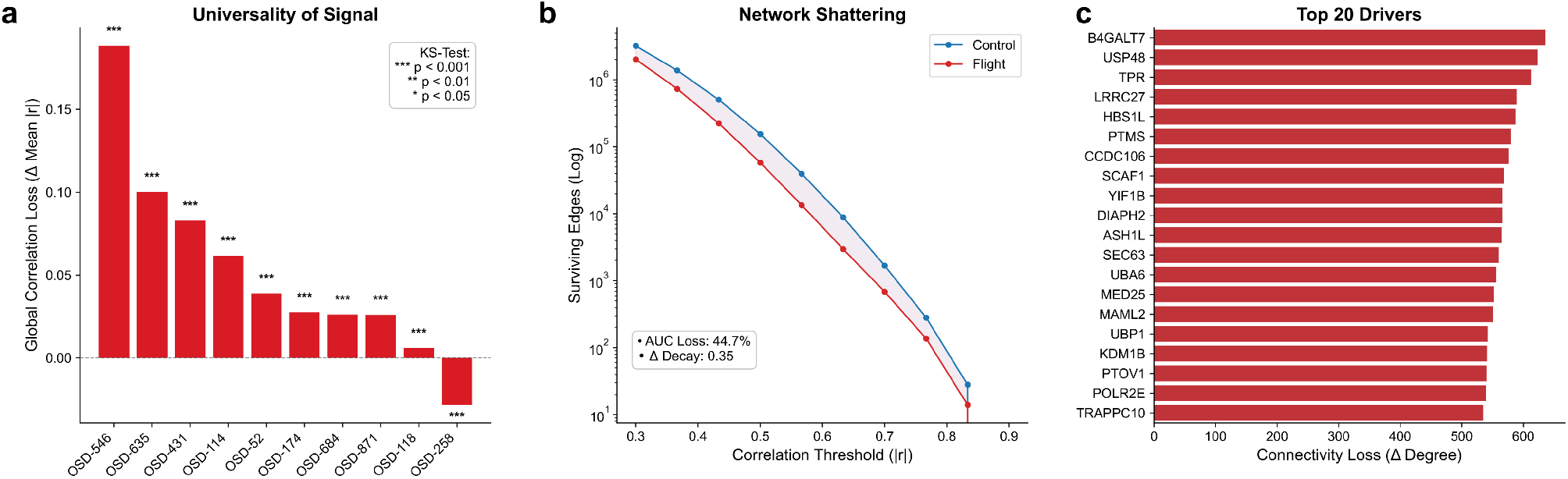
Universality and Topological Fragility. (a) Loss observed across 9/10 missions. (b) Network shattering curves. (c) Top genes driving the loss.

### Topological Fragility and Keystone Drivers

To understand the structural implications of this decoherence, we modeled the resilience of the spaceflight regulatory network via a Network Shattering analysis (Fig. 5b), wherein we mapped the decay curve of the network topology by incrementally increasing the required correlation threshold (|*r*| ) for an edge to survive. The Spaceflight network disintegrated at a significantly accelerated rate compared to Ground Control, shedding its edges prematurely. This topological fragility is quantified by a 44.7% loss in the Area Under the Curve (AUC) and a Δ Decay rate of 0.35. Physically, this indicates that the strongest, most high-confidence regulatory links are disproportionately severed in space, leaving the network structurally brittle.

To pinpoint the molecular origins of this collapse, we ranked the genome by total connectivity loss (Δ Degree) to identify specific “Keystone Drivers” (Fig. 5c). The top 20 drivers, including genes such as *B4GALT7, USP48*, and *TPR*, each lost between 500 and 650 functional regulatory partners in spaceflight. On Earth, these genes serve as highly connected central hubs maintaining network stability^47–53^. In microgravity, their sudden functional isolation likely serve as nucleation points for the broader systemic collapse observed across the genome and transcriptome.

### Integration Reveals a New Class of Regulators

Standard space omics pipelines rely on Differential Expression Analysis (DEA) to identify biomarker targets. Our DEA identified a signature of 239 canonical Differentially Expressed Genes (DEGs) meeting strict criteria (|*LFC*| *>* 0.5, FDR *p <* 0.05) which effectively clustered the samples (Fig. 6a, 6b). The DEGs identified largely recapitulate the primary findings of the original dataset investigators, heavily featuring active stress-response and DNA repair transcripts^13–22,37–46^. However, we hypothesized that focusing exclusively on transcript abundance might obscure the full scope of cellular dysregulation.

**Figure 6.**
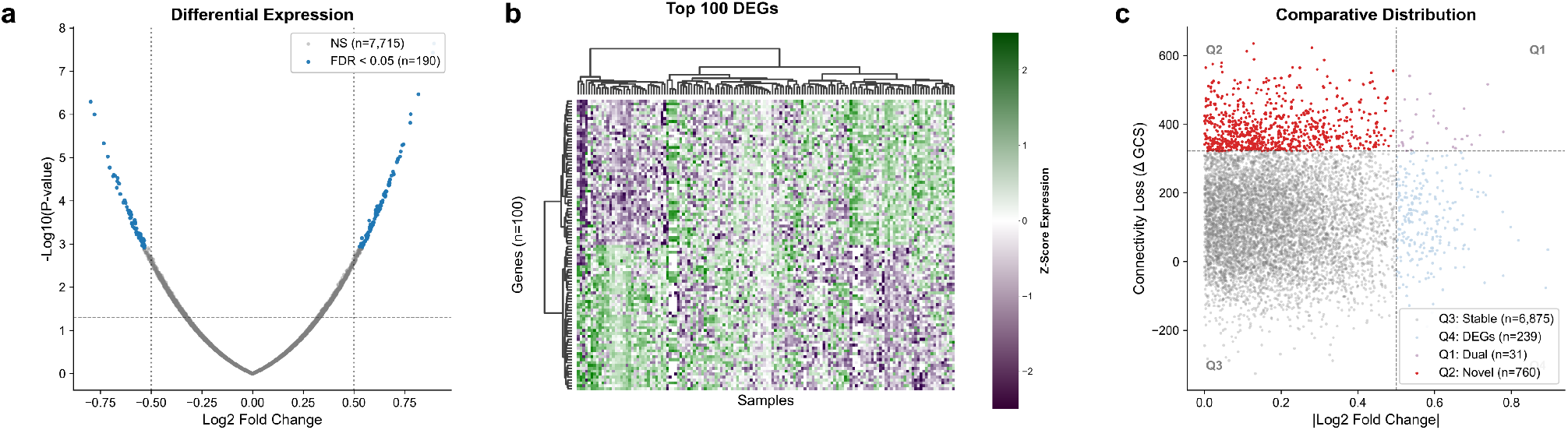
Discovery of Silent Regulators. (a) Volcano plot of canonical DEGs. (b) Heatmap of DEGs. (c) Spaceflight Quadrants identifying Silent Regulators.

Mapping the differential expression and correlation results into (Fig. 6c) revealed a decoupling of gene differential expression and correlation interactions. We identified 760 Novel Regulators (red) that exhibit high connectivity loss (Top 10%) while maintaining statistically stable expression levels (|*LFC*|*<* 0.5). We term these Silent Regulators. Interestingly, this population is more than three times larger than the population of canonical DEGs. While canonical DEGs likely represent the cell’s reactive attempt to maintain homeostasis, these genes represent the passive failure of the regulatory architecture itself, signaling hubs that are present but functionally unmoored.

### Preservation of DNA Repair vs. Metabolic Decoupling

To understand the functional consequences of this topological shift, we compared standard pathway enrichment against our novel Instability Ranking (Fig. 7). Standard over-representation analysis (Fig. 7a) highlighted DNA metabolic processes, DNA replication, and DNA repair as the most significantly perturbed pathways. This reflects a well-documented, active transcriptional response to the genomic stress and radiation environment of the ISS^54,55^.

**Figure 7.**
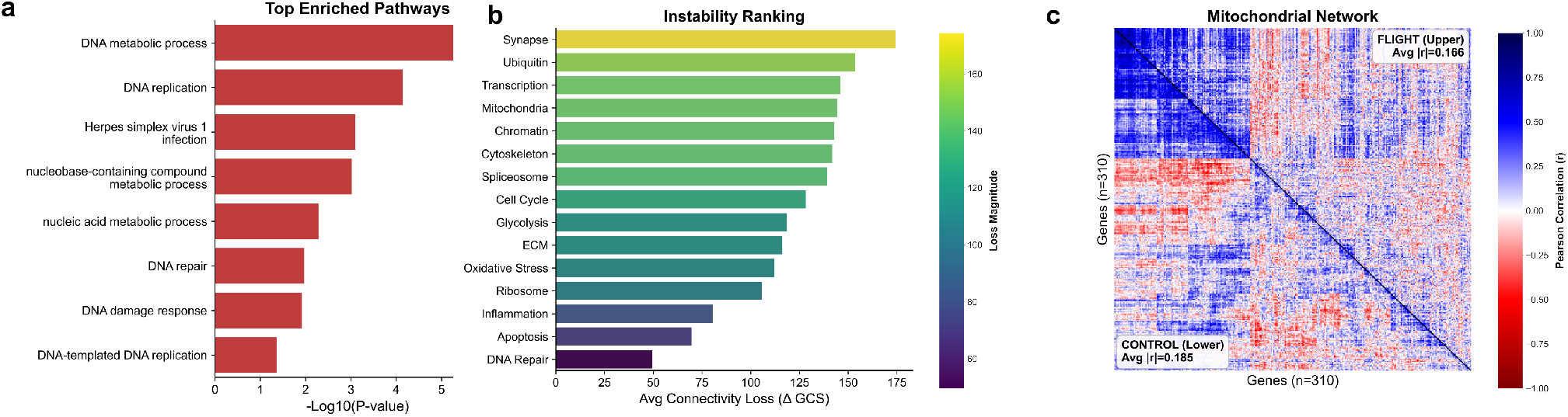
Pathway Instability Analysis. (a) Standard enrichment results. (b) Instability ranking showing mitochondrial vulnerability. (c) Targeted heatmap of mitochondrial networks.

However, our instability ranking (Fig. 7b), which evaluates topological decay rather than abundance, provided a remarkably different perspective. DNA Repair networks exhibited the lowest connectivity loss of any measured system (ΔGCS ≈ 50). This indicates that while the cell dramatically alters the expression of DNA repair genes, it expends significant effort to preserve their precise regulatory coordination, ensuring this survival-critical network does not shatter.

Conversely, systems governing higher-order physiological optimization, specifically Synaptic Signaling, Ubiquitin pathways, and Mitochondria, ranked as the most topologically unstable (ΔGCS *>* 140). We visualized this specific vulnerability by rendering a targeted correlation map of 310 core mitochondrial genes (Fig. 7c). The tight, cohesive modularity of the mitochondrial network seen on Earth (lower left) is visibly dissolved in Spaceflight (upper right), accompanied by a measurable drop in correlation strength. This metabolic decoupling suggests that the well-documented mitochondrial dysfunction experienced by astronauts is not merely a deficit in transcript availability, but a fundamental loss of stoichiometric coordination. The cell appears to execute a form of regulatory triage: allowing metabolic and synaptic networks to decohere in order to structurally protect the integrity of the genome.

### Topological Integration Maximizes Analytical Yield and Identifies Keystone Hubs

To quantify the functional importance of the newly discovered Silent Regulators, we analyzed their baseline network topology. We calculated the degree centrality (number of regulatory connections in the healthy Ground Control state) for both Canonical DEGs and Silent Regulators (Fig. 8b). Canonical DEGs exhibited a relatively low average network degree (mean ≈142 connections), typical of downstream effector genes. Conversely, Silent Regulators exhibited massively higher centrality (mean *≈* 585 connections). This mathematical distinction confirms that Silent Regulators act as keystone network hubs. Their systemic decoherence is inherently more disruptive to cellular homeostasis than the dysregulation of peripheral DEGs, further underscoring their critical importance as therapeutic targets.

**Figure 8.**
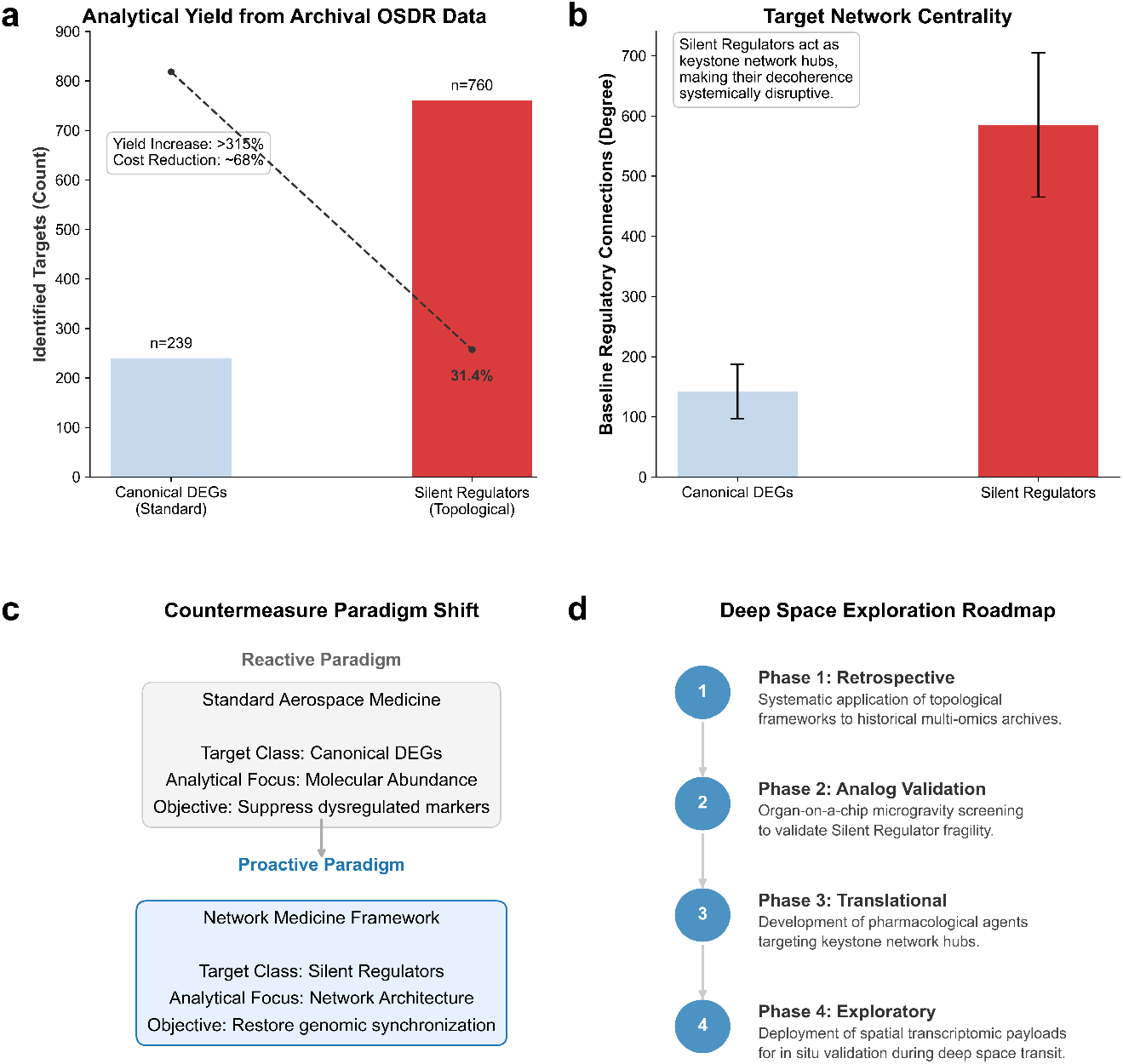
Impact of Systemic Decoherence on Aerospace Medicine and Countermeasure Development. **(a)** Analytical yield and relative cost efficiency. The integration of topological analysis identifies 760 Silent Regulators from archival data, yielding a *>* 315% increase in actionable targets compared to standard DEA (*n* = 239), effectively reducing the relative cost-per-target by ∼ 68%. **(b)** Baseline regulatory connections (network degree) for Canonical DEGs versus Silent Regulators. Silent Regulators exhibit significantly higher network centrality, confirming their role as keystone hubs whose decoherence exerts disproportionate systemic disruption. Error bars represent ± 1 standard deviation. **(c)** Proposed shift in countermeasure development. The current paradigm targets downstream DEGs to suppress symptoms, whereas the proposed proactive paradigm utilizes network topology to target Silent Regulators, aiming to restore upstream genomic synchronization. **(d)** A four-phase roadmap for integrating topological network stability into deep space exploration, spanning retrospective data mining to *in situ* exploratory validation.

Beyond the biological implications, the integration of topological algorithms provides a massive enhancement to the scientific return on investment for space omics. Biological payloads deployed to the ISS are exceptionally resource-intensive. Historically, the analytical yield of these payloads has been constrained by the limitations of standard differential expression (identifying *n* = 239 targets in this cohort). By applying our topological framework to these exact same archival datasets, we identified an additional 760 Silent Regulators, representing a *>* 315% increase in actionable biological targets (Fig. 8a). This computational re-interrogation effectively reduces the relative cost-per-target by approximately 68%, unlocking vast hidden value within historical NASA OSDR archives without requiring the deployment of novel physical payloads.

## Discussion

### Systemic Decoherence as a Thermodynamic Response to Microgravity

Our analyses identify Systemic Genome Correlation Loss as a reproducible, pan-tissue feature of the human response to spaceflight. Over the past decade, landmark investigations such as the NASA Twins Study^2^, the Space Omics and Medical Atlas^56^ and comprehensive GeneLab multi-omics analyses^1,8^ have established a foundational understanding of spaceflight adaptation. These studies successfully cataloged genomic, epigenomic, and transcriptomic shifts that characterize the spaceflight phenotype and identified consistent alterations in telomere dynamics, inflammatory cytokines, and metabolic output. Our findings build upon this foundation, demonstrating that spaceflight fundamentally alters not just the abundance of molecular features, but also the underlying regulatory architecture that coordinates them.

The data indicate a global transition from a highly structured regulatory state on Earth to a state of reduced connectivity and increased stochasticity in microgravity. This phenomenon can be understood through a thermodynamic and mechanobiological lens. Biological systems expend significant energy to maintain homeostatic order (low entropy), which manifests mathematically as tight correlations between regulatory partners. Terrestrial life evolved under the constant mechanical vector of gravity, which acts as a fundamental organizing constraint on cytoskeletal tension, fluid distribution, and nuclear architecture^57,58^. We propose that the removal of this continuous mechanical anchor reduces the energetic barrier for molecular noise, allowing genomic output to drift within a wider stochastic window. The pronounced leptokurtic shift in the global correlation distribution (Fig. 3) confirms that while the mean molecular abundance remains largely regulated, the precise couplings required for complex signal transduction are weakened. This suggests that physiological adaptation to spaceflight is intrinsically linked to a state of functional decoherence.

### Resolution of Small-Sample Artifacts via Meta-Analysis

A historical limitation in space biology has been the reliance on small-cohort studies (*N <* 6), a logistical constraint of human spaceflight. As noted in broad assessments of space omics repositories^8^, extracting consistent systemic signals from isolated datasets is statistically precarious. Mathematical theory dictates that at extremely low sample sizes, the sampling distribution of Pearson’s correlation coefficient is highly volatile; random noise can effortlessly generate spurious absolute correlations (|*r*| → 1.0), resulting in bimodal or artificially flattened distributions rather than the biologically expected Gaussian curve.

We observed this artifact in our dataset-specific supplementary analyses (Supplementary Figs. S1-S10), where individual missions with lower sample counts often exhibited dispersed correlation profiles. By aggregating 136 samples, our meta-analysis successfully converged these distributions into a mathematically robust unimodal form. This transition demonstrates that the correlation loss observed in the Spaceflight network is a genuine biological signal, distinguishable from the geometric artifacts of small sample sizes. Consequently, this study provides a mathematically stable baseline for the spaceflight transcriptome, underscoring the absolute necessity of open-science data aggregation for network-level discoveries.

### Silent Regulation: Expanding the Target Space

Current molecular countermeasures and biomarker discovery efforts predominantly target genes identified via standard differential expression analysis (DEA). For instance, established markers of the DNA damage response, such as *CDKN1A*^59,60^ and *DDB2*^61,62^, are currently utilized to monitor radiation exposure in orbit^63^. However, our integration of abundance and connectivity metrics (Fig. 6) reveals that canonical DEGs capture only a fraction of the perturbed genome. For instance, original transcriptomic analyses of these datasets reveal specific alterations in cytoskeletal and extracellular matrix expression, as well as mitochondrial dysfunction markers common to all individual datasets^13–22,37–46^. While our differential analysis confirms these reactive abundance changes, our topological framework expands this narrative significantly. This analysis identifies upstream scaffolding and metabolic genes within these same tissues that did not meet the original fold-change thresholds, yet suffered massive connectivity loss. In total, we identified a population of 760 silent regulators that exhibit stable molecular abundance but catastrophic connectivity loss.

This decoupling of abundance and topology suggests a passive failure model. For example, a transcription factor may be expressed at physiological levels, but if its co-expression with downstream targets is lost, the regulatory circuit is functionally broken. This silent population includes key drivers of cell cycle checkpoints and metabolic regulation. Because these genes do not trigger standard fold-change thresholds, they have likely been overlooked in prior candidate-gene studies. We propose that these silent regulatory genes represent the structural scaffolding of the gene regulatory network, and that their decoherence destabilizes cellular function without triggering the acute transcriptional alarms associated with inflammation or stress responses. For instance, top silent regulators identified in our analysis, such as B4GALT7 and USP48, are critical for extracellular matrix maintenance^47–49^ and DNA repair stabilization^50,51^, respectively; their functional unmooring may explain sub-clinical connective tissue and immune vulnerabilities in astronauts.

### Regulatory Triage and Metabolic Decoupling

The discrepancy between our Instability Ranking and standard pathway enrichment (Fig. 6) suggests that cells in microgravity undergo a form of preferential regulation. Standard enrichment correctly identifies DNA repair and replication as the most upregulated pathways, a known response to space radiation^64,65^. However, our topological analysis reveals that the cell actively expends energy to preserve the connectivity of these survival-critical networks. The DNA repair network exhibited the lowest instability score (ΔGCS ≈ 50), implying that upon sensing radiation damage, the cell prioritizes the stringent synchronization of existential defense mechanisms above all else.

Conversely, systems governing higher-order physiological optimization, specifically mitochondrial respiration and synaptic signaling, exhibited the highest degrees of decoherence (ΔGCS *>* 140). This targeted analysis offers a structural mechanism for hypotheses proposed by da Silveira *et al*., which established mitochondrial stress as a universal driver of spaceflight adaptation^8^. Mitochondrial function relies on the precise, stoichiometric assembly of the Electron Transport Chain (ETC), necessitating flawless coordination between nuclear and mitochondrial genomic outputs. Our data suggests that mitochondrial dysfunction in space is not exclusively a failure of biogenesis, but a fundamental failure of assembly. When the correlations between these subunits dissolve (Fig. 7c), the ETC cannot assemble efficiently, which may lead to reactive oxygen species (ROS) leakage and energetic deficits, even if the constituent transcript parts remain abundant^66^. The cell functionally sacrifices metabolic efficiency to conserve the regulatory energy required to maintain genomic integrity.

### Topological Fragility and Clinical Implications

Network analysis (Fig. 5b) also highlights a latent clinical risk. While the spaceflight regulatory network remains functional under steady-state conditions, it retains significantly fewer redundant high-confidence edges than the ground control network. In network theory, connectivity confers resilience against perturbations^9^. The rapid disintegration of the spaceflight network under increasing stringency thresholds implies a reduced buffering capacity.

This topological fragility suggests that astronauts may be hypersensitive to subsequent environmental challenges. Clinical observations from prolonged ISS missions frequently document phenomena such as latent viral reactivation (e.g., Epstein-Barr virus, Varicella-Zoster virus) despite seemingly normal baseline leukocyte counts^67,68^. Our findings offer a topological basis for this disconnect: the immune and stress-response networks, while physically present in abundance, are loosely coupled and brittle. Consequently, while an astronaut’s physiology is nominally compensated, the weakened regulatory web leaves them highly vulnerable to acute secondary stressors such as unexpected pathogens, extreme stress/trauma, or radiation events. A biological network operating without a structural safety margin may collapse catastrophically under external loads that would be easily buffered on Earth.

### Limitations and Future Directions

While this study provides the most comprehensive topological view of the spaceflight genome and transcriptome to date, it relies on nucleic acid profiles as a proxy for functional biological networks. While mRNA and genomic co-expression are robust predictors of regulatory topology, post-transcriptional buffering (e.g., phosphorylation cascades, translational regulation) could theoretically mitigate some of this decoherence at the final proteome level. Future multi-omics inquiries utilizing matched multi-omics and integrative studies are essential to determine the extent to which this regulatory entropy propagates across omic layers. Furthermore, while batch-specific normalization successfully harmonized diverse platforms, the reliance on bulk tissue datasets inherently averages out cell-type-specific responses. Single-cell covariance analysis represents the logical next frontier to ascertain whether decoherence arises from intracellular de-synchronization or from increased phenotypic heterogeneity between individual cells within a tissue.

### Moving Forward

As humanity crosses the threshold from LEO operations to sustained interplanetary exploration via the Artemis and Martian programs, understanding the absolute limits of human physiological resilience is paramount. Our analysis demonstrates that adaptation to the spaceflight environment cannot be fully comprehended through the traditional lens of differential molecular abundance alone. By successfully harmonizing highly heterogeneous transcriptomic and genomic datasets into a unified topological framework, we have identified Systemic Genome Correlation Loss as a central, universal biosignature of spaceflight. This discovery fundamentally shifts the paradigm of space omics: the primary threat to cellular homeostasis in microgravity is not merely the aberrant expression of specific genes, but the entropic dissolution of the relationships between them.

We have shown that the spaceflight expression profile regresses towards a state of increased stochastic noise, and that this topological decay appears to be driven by a previously unrecognized class of silent regulator genes. Because these hub genes maintain stable baseline expression levels, they systematically evade standard statistical detection, yet their functional detachment from downstream targets nucleates widespread systemic instability. Furthermore, our findings suggest that this decoherence is not entirely random, but reflects a calculated cellular triage. By expending regulatory energy to preserve existential DNA repair networks while allowing highly complex metabolic and synaptic systems to shatter, the human body fundamentally re-prioritizes immediate cellular survival over higher-order physiological optimization. This metabolic decoupling provides a unifying molecular explanation for the physiological symptoms experienced by astronauts.

Consequently, future spaceflight medicine must evolve beyond symptom management and targeted pathway inhibitors. The discovery of systemic decoherence expands possible countermeasure development to incorporating network stabilizers, interventions specifically engineered to reinforce regulatory coupling, restore mitochondrial stoichiometry, and re-anchor the genome’s structural fidelity. Furthermore, advancing this framework via single-cell spatial omics and matched proteomic covariance mapping will be essential to precisely locate where these network fractures originate. Monitoring and mitigating spaceflight-induced correlation loss serves as an additional metric to ensure that humanity not only survives the extreme environments we will be exposed to in our future in space, but maintain the physiological coherence required to thrive within them.

## Acknowledgments

We gratefully acknowledge the NASA Open Science Data Repository (OSDR) and the GeneLab project for their commitment to democratizing access to spaceflight omics data. Open science initiatives are essential for maximizing the scientific return of spaceflight missions and enabling the discovery of systemic biosignatures that transcend individual experiments. We thank the original principal investigators and research teams of the 10 datasets analyzed in this study for their dedication to conducting rigorous science in the challenging environment of space.

## Data Availability

The datasets analyzed during the current study are publicly available in the NASA Open Science Data Repository (OSDR) under the accession numbers: OSD-52^13^, OSD-114^14^, OSD-118^15^, OSD-174^16^, OSD-258^17^, OSD-431^18^, OSD-546^19^, OSD-635^20^, OSD-684^21^, and OSD-871^22^ (https://osdr.nasa.gov).

## Code Availability

The complete Python analysis pipeline is available in the GitHub repository at github.com/AnuragSakharkar/Systemic-Genome-Correlation-Loss.

## Authorship Contributions

A.S. and A.J.B. conceived the study, designed the analytical framework, and performed the primary data analysis. A.J.B., C.E.M., C.T., J.Y., and K.E.L. supervised the conceptualization and execution of the research. L.S. and S.C. provided critical contributions to all aspects of the bioinformatic analyses and data harmonization. A.S. and A.J.B. co-wrote the initial draft of the manuscript. All authors engaged in reviewing, editing, and refining the manuscript, and all authors approved the final submitted version.

