## Supplemental document for "Systemic Genome Correlation Loss as a Central Characteristic of Spaceflight"

Anurag Sakharkar et al.

**S2 Global Data Normalization Quality Control** **3**

**S3 Supplementary Figures: Individual Mission Analysis** **4**

### S1 Computational Pipeline Architecture

The bioinformatic analysis presented in this study was executed via a modular Python pipeline designed for high-dimensional topological analysis. The codebase is structured to ensure reproducibility, statistical rigor, and visual standardization across heterogeneous transcriptomic and genomic datasets. The workflow proceeds through four distinct phases:

#### S1.1 Data Ingestion and Harmonization

The pipeline begins by ingesting raw transcriptomic/genomic count matrices or normalized intensity files from the `data/00_raw/` directory.

- **Strict Intersection:** Gene identifiers are mapped to HGNC symbols using a static BioMart reference. Only genes detected across all 10 datasets are retained to prevent imputation artifacts.
- **Batch-Specific Normalization:** A custom Z-score normalization algorithm is applied independently to each dataset (Batch) before merging. This ensures that the statistical moments (mean and variance) are aligned across platforms (Microarray vs. RNA-Seq) while preserving the relative biological signal between Control and Flight samples within each mission.

#### S1.2 Metric Calculation (Parallelized)

Given the computational cost of calculating genome-wide correlation matrices ($N \approx 15,000$ genes, $> 10^8$ pairwise interactions), the pipeline utilizes the `joblib` library with a `threading` backend to leverage multi-core CPU architectures.

- **Global Connectivity Score (GCS):** The pipeline computes the sum of absolute Pearson correlation coefficients for every gene against the entire genome.
- **Decoherence Metric:** Defined as $\Delta GCS = GCS_{Control} - GCS_{Flight}$.

#### S1.3 Statistical Validation Framework

To differentiate biological signal from noise, the pipeline executes three rigorous validation routines:

1. **Saturation Analysis:** A bootstrapping module iteratively subsamples the dataset (from $N = 4$ to $N_{max}$) to determine the stability threshold, a sample size at which the signal-to-noise ratio drops below 0.25 and the mean loss converges.
2. **Permutation Testing:** A parallelized module shuffles sample labels 2,500 times to generate a null distribution, calculating empirical p-values for the observed correlation loss.
3. **Variance Independence:** A regression module stratifies genes by variance quantiles to verify that connectivity loss is not an artifact of low-expression noise.

#### S1.4 Visual Assembly System

Figures are generated to ensure uniform styling, scaling, and font sizes across the manuscript.

- **Discrete Generation:** Each subplot is generated as a standalone high-resolution (600 DPI) image using strict geometric constraints (e.g., $6 \times 6$ inch squares).
- **Panel Stitching:** `matplotlib.gridspec` is used to stitch these discrete images into the final composite figures. This approach prevents the distortion of text labels and aspect ratios often seen when resizing complex vector graphics.

#### S2 Global Data Normalization Quality Control

To ensure that the Batch-Specific Z-score normalization successfully aligned the heterogeneous datasets without introducing artifacts or erasing biological variance, we performed a global quality control assessment of the standardized expression matrix.

Figure S1 displays the distribution of gene expression Z-scores for 25 randomly selected samples across the 10 missions. The consistent centering of the medians at zero and the alignment of interquartile ranges (IQR) across the cohort demonstrate that the normalization effectively mapped disparate measurement scales (from both Microarray and RNA-Seq) into a common dimensionless space. This alignment is a prerequisite for the high-dimensional correlation analyses performed in this study, as it ensures that the calculated Pearson coefficients ($r$) reflect shared biological covariance rather than technical baselines.

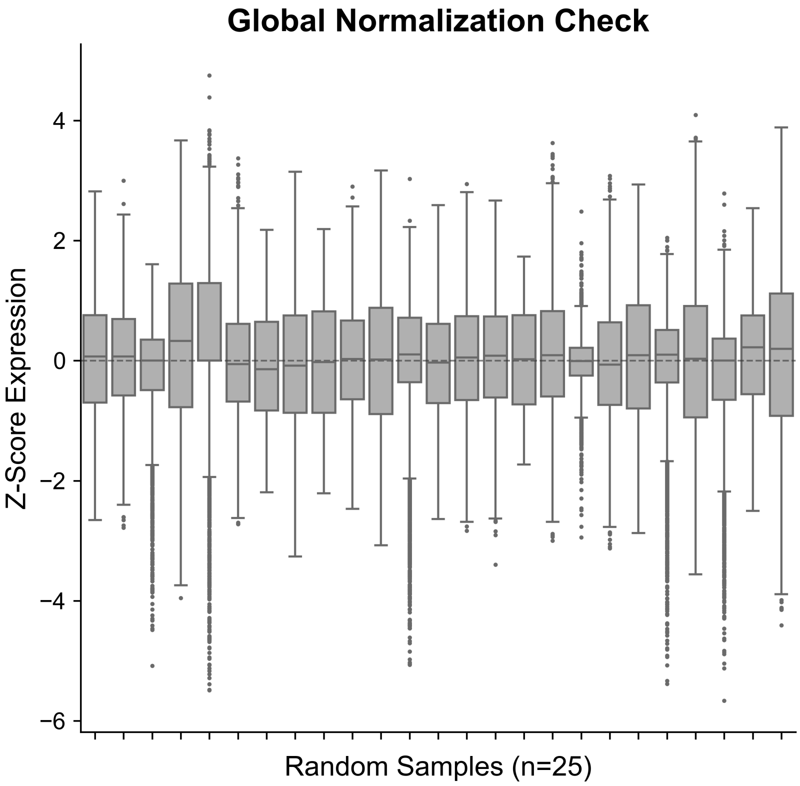

Figure S1: **Global Normalization Check.** Boxplot showing the distribution of normalized expression values (Z-scores) for 25 randomly selected samples from the integrated meta-cohort. Medians are centered at 0 (red dashed line) with consistent variance across samples, confirming successful batch-effect mitigation across the 10 constituent datasets.

#### S3 Supplementary Figures: Individual Mission Analysis

To verify the universality of Systemic Genome Correlation Loss, we performed a complete independent analysis for each of the 10 datasets included in the meta-analysis. For each mission, we generated:

- **(a) Mirror Plot:** Visualizes the structural erosion of gene-gene correlations (Control vs. Flight).
- **(b) Distribution Shift:** Quantifies the regression of correlation coefficients toward zero (Leptokurtosis).
- **(c) Network Shattering:** Demonstrates the accelerated decay of network edges under increasing stringency thresholds.
- **(d) Normalization QC:** Violin plots of Z-score distributions for Control and Flight samples, confirming effective centering and variance stabilization.

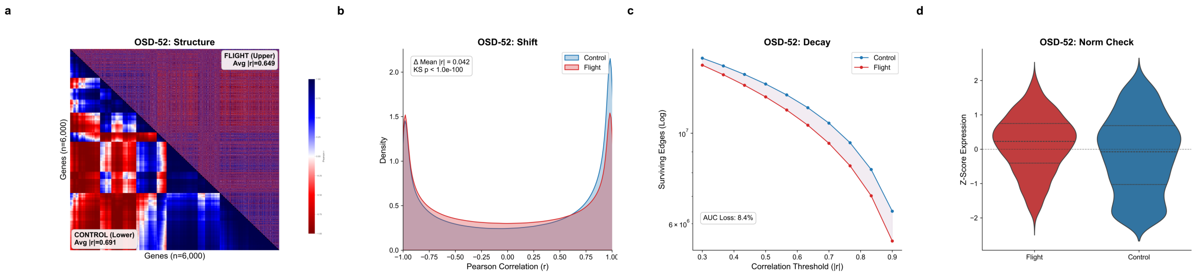

Figure S2: **OSD-52 (Endothelial Cells, 10 Days).** (a) Mirror plot showing loss of structural definition in Flight (upper triangle). (b) Correlation distribution shows a significant shift toward zero. (c) Flight network (red) shatters faster than Control (blue). (d) Post-normalization QC confirms aligned distributions.

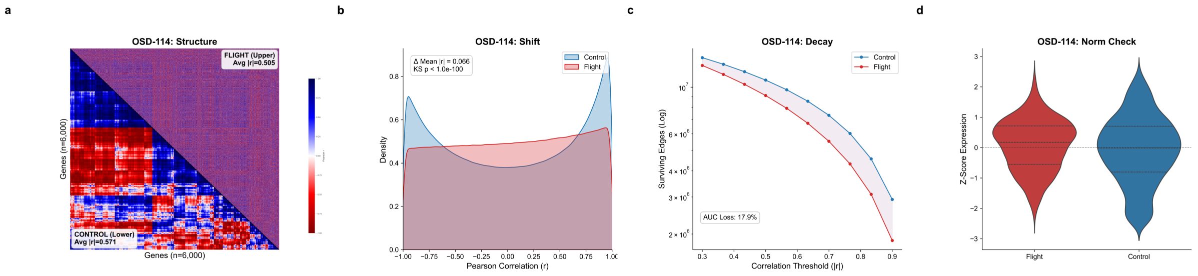

Figure S3: **OSD-114 (Skin, 3 Days).** (a) Mirror plot visualization. (b) Distribution shift analysis. (c) Network shattering curve indicating topological fragility in spaceflight. (d) Post-normalization QC.

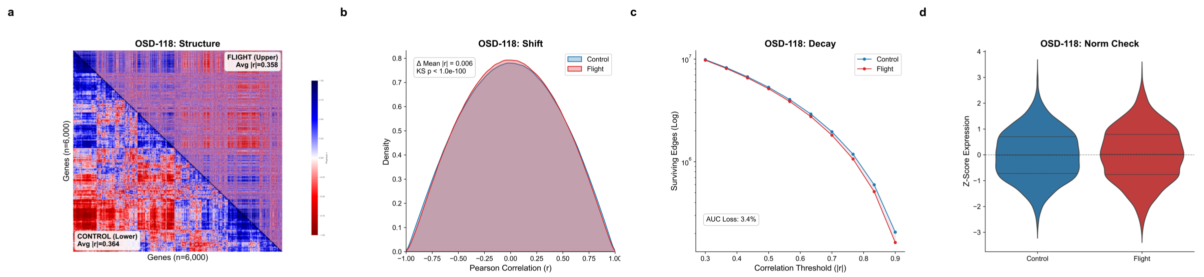

Figure S4: **OSD-118 (Skin, 14 Days).** (a) Mirror plot visualization. (b) Distribution shift analysis. (c) Network shattering curve. (d) Post-normalization QC.

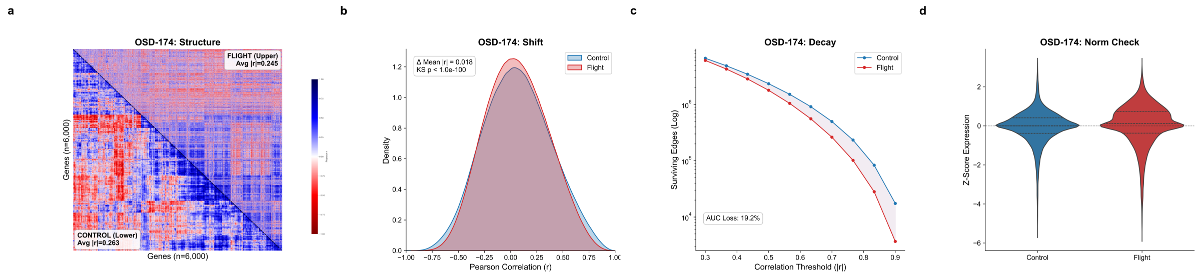

Figure S5: **OSD-174 (Hair Follicle, 37 Days).** (a) Mirror plot visualization. (b) Distribution shift analysis. (c) Network shattering curve. (d) Post-normalization QC.

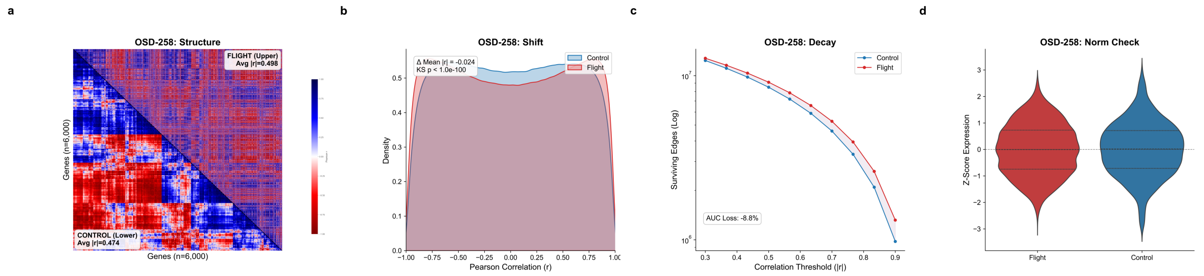

Figure S6: **OSD-258 (Cardiomyocytes, 30 Days).** (a) Mirror plot visualization. (b) Distribution shift analysis. (c) Network shattering curve. (d) Post-normalization QC.

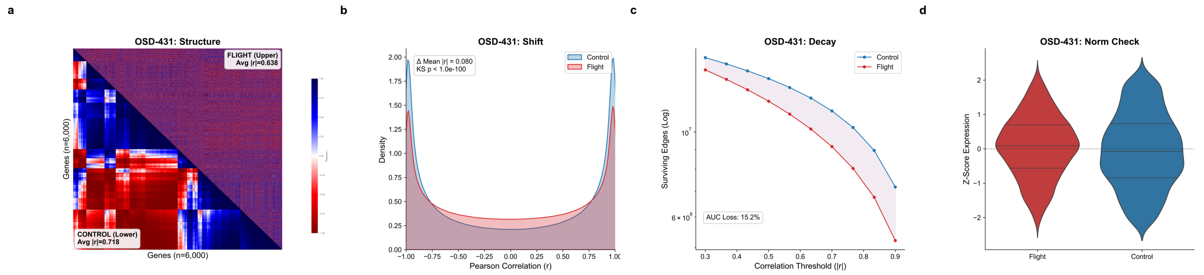

Figure S7: **OSD-431 (Skin, 160 Days).** (a) Mirror plot visualization. (b) Distribution shift analysis. (c) Network shattering curve. (d) Post-normalization QC.

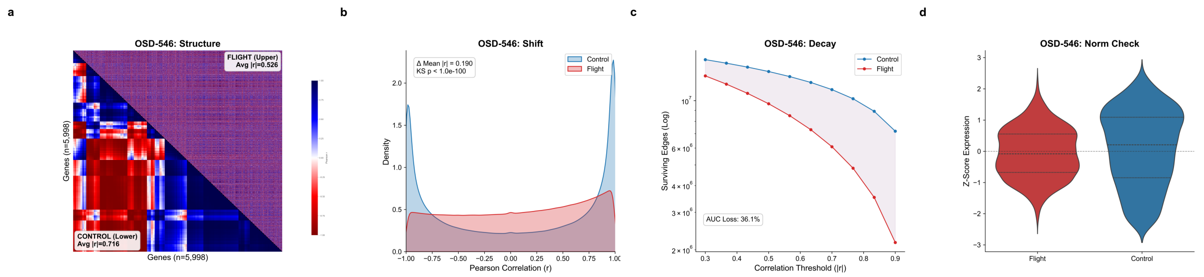

Figure S8: **OSD-546 (Bone Marrow, 14 Days).** (a) Mirror plot visualization. (b) Distribution shift analysis. (c) Network shattering curve. (d) Post-normalization QC.

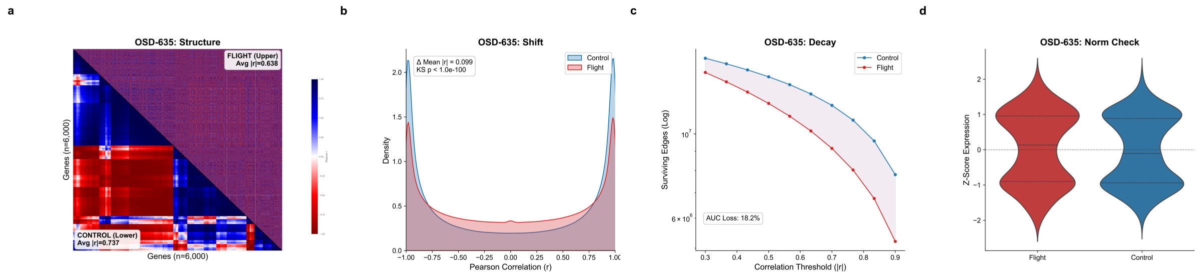

Figure S9: **OSD-635 (Smooth Muscle, 3 Days).** (a) Mirror plot visualization. (b) Distribution shift analysis. (c) Network shattering curve. (d) Post-normalization QC.

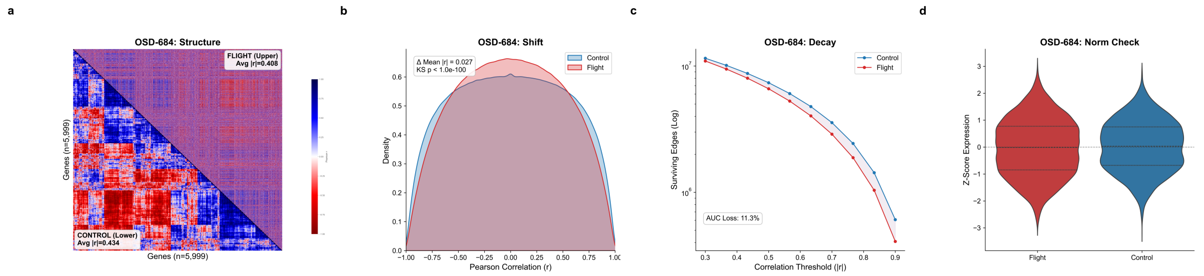

Figure S10: **OSD-684 (Myoblasts, 5 Days).** (a) Mirror plot visualization. (b) Distribution shift analysis. (c) Network shattering curve. (d) Post-normalization QC.

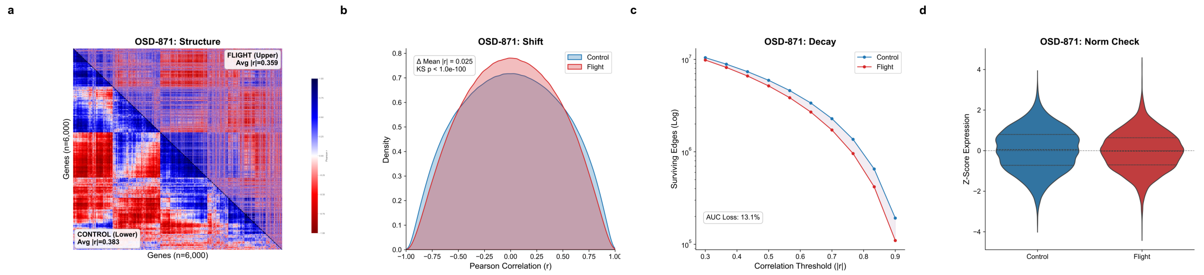

Figure S11: **OSD-871 (iPSC, 30 Days).** (a) Mirror plot visualization. (b) Distribution shift analysis. (c) Network shattering curve. (d) Post-normalization QC.
